# Employing connectome-based models to predict working memory in multiple sclerosis

**DOI:** 10.1101/2021.03.01.432930

**Authors:** Heena R. Manglani, Stephanie Fountain-Zaragoza, Anita Shankar, Jacqueline A. Nicholas, Ruchika Shaurya Prakash

**Author notes:** <u>Corresponding Author:</u> Ruchika Shaurya Prakash, Department of Psychology, The Ohio State University, 1835 Neil Ave Columbus, OH 43210.

## Abstract

**Background:** Individuals with multiple sclerosis (MS) are vulnerable to deficits in working memory, and the search for neural correlates of working memory in circumscribed areas has yielded inconclusive findings. Given the widespread neural alterations observed in MS, predictive modeling approaches that capitalize on whole-brain connectivity may better capture individual-level working memory in this population.

**Methods:** Here, we applied connectome-based predictive modeling to functional MRI data from working memory tasks in two independent samples with relapsing-remitting MS. In the internal validation sample (*n*_internal_ = 36), functional connectivity data were used to train a model through cross-validation to predict accuracy on the Paced Visual Serial Addition Test, a gold-standard measure of working memory in MS. We then tested its ability to predict performance on the N-back working memory task in the external validation sample (*n*_external_ = 36).

**Results:** The resulting model successfully predicted working memory in the internal validation sample but did not extend to the external sample. We also tested the generalizability of an existing model of working memory derived in healthy young adults to people with MS. It showed successful prediction in both MS samples, demonstrating its translational potential. We qualitatively explored differences between the healthy and MS models in intra- and inter-network connectivity amongst canonical networks.

**Discussion:** These findings suggest that connectome-based predictive models derived in people with MS may have limited generalizability. Instead, models identified in healthy individuals may offer superior generalizability to clinical samples, such as MS, and may serve as more useful targets for intervention.

**Impact Statement:** Working memory deficits in people with multiple sclerosis have important consequence for employment, leisure, and daily living activities. Identifying a functional connectivity-based marker that accurately captures individual differences in working memory may offer a useful target for cognitive rehabilitation. Manglani et al. demonstrate machine learning can be applied to whole-brain functional connectivity data to identify networks that predict individual-level working memory in people with multiple sclerosis. However, existing network-based models of working memory derived in healthy adults outperform those identified in multiple sclerosis, suggesting translational potential of brain networks derived in large, healthy samples for predicting cognition in multiple sclerosis.

## Introduction

Multiple sclerosis (MS) is the most common demyelinating disease of the central nervous system, driven by chronic inflammatory, neurodegenerative processes. Nearly one million individuals in the United States and 2.5 million worldwide are living with the disease (Wallin *et al*., 2019), of whom, at least half will exhibit cognitive deficits during their clinical course (Chiaravalloti and DeLuca, 2008). Among the cognitive sequelae which intensify personal and economic burden of the disease, working memory dysfunction is notable (Macías Islas and Ciampi, 2019). Working memory (WM) is the brain’s ability to temporarily store, prioritize, and actively manipulate transitory information. In the constant competition between relevant and irrelevant data, deficits in maintaining target information can result in slower or inaccurate processing, thereby causing demonstrable difficulties in daily tasks that rely on WM (Cabeza *et al*., 2016). WM deficits in people with MS (PwMS) are associated with functional ramifications including lower work engagement and unemployment (Macías Islas and Ciampi, 2019; Nicholas *et al*., 2019). Given the neuropathology characteristic of MS, various neuroimaging investigations have sought to understand how neural processes supporting WM may be impacted by the disease.

Existing studies of WM in MS have predominantly focused on *a priori* regions and/or networks of interest, yielding evidence limited to select brain areas. Some functional MRI (fMRI) studies show that during WM tasks, PwMS demonstrate greater activity of prefrontal regions than healthy controls (Forn *et al*., 2006; Forn *et al*., 2007; Hillary *et al*., 2003). However, other research indicates that activity patterns depend on individual WM capacity; relative to healthy controls, unimpaired PwMS show similar activity patterns, whereas WM-impaired individuals exhibit greater recruitment of frontal and parietal regions (Chiaravalloti *et al*., 2005). Neural recruitment also appears to depend on task demand. Under low WM demand, PwMS activate prefrontal and parietal areas to a greater degree than healthy controls (Mainero *et al*., 2004; Sweet *et al*., 2006; Colorado *et al*., 2012), suggesting a compensatory response. In contrast, when faced with high WM demand, PwMS exhibit a pattern that is thought to be maladaptive (Giorgio *et al*., 2015; Vacchi *et al*., 2017), characterized by reduced recruitment of core prefrontal and parietal regions, and greater activation of areas outside of typical WM circuitry (Wishart *et al*., 2004; Sweet *et al*., 2006; Vacchi *et al*., 2017).

Recent fMRI studies on the relationship between WM and communication among intrinsic functional networks reveal diffuse connectivity changes in MS. Relative to healthy controls, PwMS show increased functional connectivity in sensorimotor and cognitive control networks at rest (Giorgio *et al*., 2015). Additionally, there is evidence that disease progression may be associated with more widespread alterations in communication between disparate brain areas. Compared to healthy controls and unimpaired PwMS, cognitively impaired PwMS demonstrate increased connectivity between both the default mode and the frontoparietal networks with other brain networks (Meijer *et al*., 2017). However, across studies, differences in regional activity or connectivity between select networks does not adequately distinguish individuals with worse clinical course (Vacchi *et al*., 2017). This may be due to the heterogeneous disease presentation, whereby pathology and neural dysfunction impacts a wide swath of cortical and subcortical regions (Rovaris *et al*., 2000).

The predominant use of correlational methods for discovering associations between brain activity and WM have proven insufficient for converging upon a single reliable signature of WM in MS. Previous studies are limited in that they: 1) are restricted to specific areas/networks of the brain; 2) rely on group-level analysis, yielding sample-specific findings; 3) ignore rich individual-level variability; and 4) do not test generalizability of findings in novel samples. Predictive modeling methods that capitalize on whole-brain functional connectivity offer a promising approach for identifying a WM neural signature in MS. In fact, a growing literature indicates that individual differences in distributed patterns of functional connectivity can be harnessed to identify neuromarkers that predict cognition (Rosenberg *et al*., 2017; Greene *et al*., 2018; Barron *et al*., 2019). Given the heterogeneous and widespread neural alterations observed in MS, broadening the search field for brain-behavior relationships may help identify a comprehensive network-based model of WM in MS.

Connectome-based predictive modeling (CPM) is a data-driven technique for identifying whole-brain functional networks that can be used to make predictions about cognitive performance in previously unseen individuals (Shen *et al*., 2017). CPM has been used to successfully predict a wide range of cognitive measures, including sustained attention (Rosenberg *et al*., 2016), reaction time and executive control (Rosenberg *et al*., 2017), fluid intelligence (Finn *et al*., 2015; Greene *et al*., 2018), and WM (Avery *et al*., 2019) in healthy young-adult samples as well as cognitive function in normal aging and neurodegenerative disease (Lin *et al*., 2018, Avery *et al*., 2019; Fountain-Zaragoza *et al*., 2019), suggesting successful clinical translation. Of particular relevance, Avery and colleagues (2019) used CPM to identify functional networks during a WM task that predicted performance in healthy adults. This working memory CPM (wmCPM) successfully predicted memory performance in an independent, heterogeneous sample of older adults with preserved cognition, amnestic mild cognitive impairment, and Alzheimer’s disease. The demonstrated ability of CPM to yield network-based models that predict individual differences in WM and generalize to clinical populations suggests utility of this approach for predicting WM in PwMS.

The primary aims of this study were to: 1) derive a connectome-based predictive model of WM in MS (MS-wmCPM), and (2) test the generalizability of this model to an independent sample of PwMS and an alternative task of WM. A secondary aim was to apply an existing wmCPM derived in healthy young adults (Avery *et al*., 2019) to both samples of PwMS to assess the translational potential. Additionally, we explored differences between the wmCPM and MS-wmCPM in topology and involvement of canonical networks.

## Materials and methods

### Participants

Two independent samples were used for deriving the WM model (internal validation sample) and testing prediction in an independent group (external validation sample). Individuals with relapsing remitting MS were recruited from Columbus, Ohio and the surrounding areas between 2014-2016 (internal sample) and 2017-2021 (external sample). Eligibility requirements for both studies included: normal or corrected vision ≥ 20/40, without cognitive impairment indicated by a score >23 on the Mini-Mental Status Exam (Folstein *et al*., 1975; range_internal_ = 25-30, range_external_ = 24-30), without significant depressive symptoms assessed by a Beck Depression Inventory score ≤ 19 (Beck *et al*., 1996), right-handedness, ability to walk without aid for 100 meters indicated by a self-reported Expanded Disability Status Scale score between 0-5.5 (Kurtzke, 1983), absence of comorbid neurologic or psychiatric disorders, and free of relapse and corticosteroid use in the prior 30 days.

Data for the internal validation sample were initially acquired as part of a cross-sectional study assessing the relationship between physical activity and graph theoretical metrics. Data for the external validation sample were drawn from baseline (prior to randomization) assessments of a randomized controlled trial (Clinicaltrials.gov: NCT03244696) comparing the effects of physical activity with hydration on cognition. In both samples, participants were excluded from further analysis if they demonstrated excessive head motion (see “Motion controls” in the Supplementary Materials) or were deemed by the collaborating neurologist to have incidental findings on structural imaging atypical of MS pathology. Ultimately, the internal validation sample included 36 participants for model derivation (29 women, aged 30-58 years, *M*_*age*_ = 45.5, mean disease duration = 10.8 years) and the external validation sample included 36 participants (28 women, ages 31-58, *M*_*age*_ = 45.3, mean disease duration = 9.99 years) for testing model generalizability. For both studies, informed consent was collected in accordance with The Ohio State University Institutional Review Board. Networks from Avery et al. (2019) were generated on 502 participants (274 women, aged 22-35 years, *M*_*age*_ = 28, *SD* = 3.6 years).

### Paradigms

The internal validation sample completed the Paced Visual Serial Addition Test (PVSAT; Fos *et al*., 2000) during scanning. Participants were asked to add two numbers presented either consecutively (serial condition), or simultaneously (math condition), and indicate by button press whether the sum was > or <10. The task was divided into four alternating serial and math blocks with 15 trials per block for a total duration of ∼14 min. The dependent variable of interest was average accuracy across all serial blocks.

The external validation sample completed the N-back task from the Human Connectome Project, described in detail elsewhere (Van Essen *et al*., 2012; Barch *et al*., 2013), during scanning. Participants were presented with pictures of places, tools, faces, and body parts and asked to indicate by button press whether each subsequent image matched the target (0-back condition), or the image two previous to the current image (2-back condition). The task included two runs, each with eight alternating blocks (four 0-back, four 2-back), and 10 trials per block for a total duration of ∼15 min. Accuracy in the 2-back condition served as the dependent variable of interest.

### Network Construction

Data for both studies were collected at The Ohio State University Center for Cognitive and Behavioral Brain Imaging on a 3T Siemens scanner with a 32-channel head coil. Acquisition parameters for both studies (Supplementary Table 1) and additional processing details are provided in the Supplementary materials.

#### Whole-brain functional connectivity

Preprocessing steps included brain extraction and lesion filling of the T1w image; motion and distortion correction, slice timing correction for non-multiband data, brain extraction, spatial smoothing of the 4D data; and registration of 4D data to the participant’s T1w lesion-filled image. Nuisance variables (average white matter, gray matter, cerebrospinal fluid, and global signal) were regressed from the 4D data and the residual data were advanced to the Graph Theory General Linear Model (GTG) toolbox (Spielberg *et al*., 2015) for estimation of a whole-brain functional connectivity matrix for each participant, comprised of edges (i.e., functional connections) between all nodes (i.e., brain regions). Brains were parcellated using the 268-node Shen functional atlas spanning cortical, subcortical, and cerebellar regions (Shen *et al*., 2013). Mean activity in each node was calculated as the average blood-oxygen level-dependent signal across time within all constituent voxels. Functional connectivity was indexed by the Pearson correlation between the mean timecourse of every pair of nodes. All values were then Fisher *r*-to-*z* transformed. Sixteen nodes missing sufficient coverage (<5 voxels) in at least one participant (seven in each right and left cerebellum, one in the right brainstem, and one in the left temporal lobe) were excluded from internal validation analysis, resulting in a 252 x 252 symmetrical connectivity matrix for each participant.

#### Internal model fitting

Connectome-based predictive modeling (Shen *et al*., 2017) was implemented using custom MATLAB scripts (available here: https://www.nitrc.org/projects/bioimagesuite/) to identify functional networks that were predictive of PVSAT scores in the internal validation sample. Within each of the 36 rounds of leave-one-out cross validation (LOOCV), steps included feature selection, model building, and model-based prediction. Feature selection involved conducting a Spearman’s rank correlation between each edge and PVSAT performance across the training set. Correlations were thresholded at *p* < .01 and divided into edges that were positively associated with performance (high-WM network) and negatively associated with performance (low-WM network). Network strengths were computed as the average connectivity among edges included in the high- and low-WM networks. Combined WM network strength was computed as the difference between strengths in the high- and low-WM networks. Model building involved conducting three linear regressions to separately fit network strengths (high, low, and combined) to WM scores in the training set. Last, network strengths for the left-out test participant were computed and entered into each model to generate predicted WM scores based on the low, high, and combined models. These steps were repeated until all participants had served as the left-out sample once. Model performance was assessed as the Spearman’s rank correlation between predicted and observed WM scores. To construct final high- and low-WM models (MS-wmCPM), edges found in every round of cross validation were retained, and a general linear model was built to fit strength in those edges to PVSAT scores in the full internal sample. In addition to model derivation using linear regression, we tested our model using ridge regression. This technique includes a regularization *λ* function, which may shrink regression coefficients (Hoerl and Kennard, 2000) to account of multicollinearity among edges, counter overfitting in small samples, and improve model generalizability to unseen samples (Cui and Gong, 2018). Permutation testing, involving random shuffling of brain-behavior relationships, was conducted across 1000 iterations to generate permuted *p*-values.

#### External validation

As the gold-standard assessment of predictive power (Scheinost *et al*., 2019), the MS-wmCPM models were applied to the previously unseen, external validation sample. As described above, we calculated whole-brain functional connectivity during the N-back task for each participant and then calculated high-, low-, and combined MS-wmCPM network strengths for each participant. Model performance, or the correspondence between network strength and observed N-back scores, was assessed using multiple measures of predictive power outlined in the “Statistical Analysis” section below.

#### Comparing Predictive Ability

The predictive abilities of the MS-wmCPM and WM networks derived in healthy young adults (wmCPM; Avery *et al*., 2019) were compared. For both samples of PwMS, each participant’s functional connectivity in the included wmCPM edges was averaged to compute high, low, and combined network strengths. All correlations were Fisher *r*-to-*z* transformed and the Dunn and Clark’s *z* test (1971) was computed using the statistical comparison package cocor (Diedenhofen and Musch, 2015), which compares values from the same (i.e., dependent) sample with overlapping (i.e., observed WM) scores.

#### Functional Anatomy of WM Networks

To account for the size of each macroscale network, including medial frontal, frontoparietal, default mode, motor, visual I, visual II, visual association, salience, subcortical, and cerebellar, an index of contribution was calculated using the formula from Greene and colleagues (2018). Each edge between nodes *i* and *j* was assigned to the respective pair of canonical networks, *A* and *B*. In this contribution index, *M*_*A,B*_ denotes the number of edges between those canonical networks in the model, and *M*_*tot*_ is the total number of edges in the MS-wmCPM. *E*_*A,B*_ is the number of edges between networks *A* and *B* in the whole brain, and *E*_*tot*_ is the total number of edges in the whole brain. A relative contribution value > 1 indicated a disproportionately greater contribution of that network pair to the model.

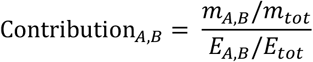

### Statistical Analysis

Statistical analyses were performed using SPSS 26.0 (SPSS, Chicago, IL, USA) and MATLAB 2019a (MathWorks, Natick, MA, USA). The Shapiro-Wilk and χ^2^ tests were used to assess normality of all continuous and categorical variables, respectively. Independent samples t-tests and Mann Whitney *U* tests assessed differences between the samples on demographic and clinical characteristics. Two-tailed statistical tests with an alpha of .05 were used to determine significance. As predicted and observed values are not expected to lie on the same scale and only correspond in relative terms, model performance was assessed using Spearman’s rank correlation (*r*_s_) between observed WM scores and summary strength of the high, low, and combined networks. As recommended by Scheinost *et al*. (2019), additional indicators of model performance included mean squared error (MSE) and prediction *R*^2^ (hereinafter referred to as *R*^2^), calculated as 1 minus the normalized mean squared error between observed and predicted values.

### Data availability

The study data and code can be made available upon request from the corresponding author.

## Results

### Demographics, Clinical Characteristics, and Behavioral Performance

Table 1 summarizes demographic, clinical, and behavioral data for both samples. There was a significant difference in total lesion volume between the samples (*U* = 308, *p* < .001). There were no significant differences between the samples in any of the other demographic or clinical variables (all *p*s > .38).

**Table 1.**
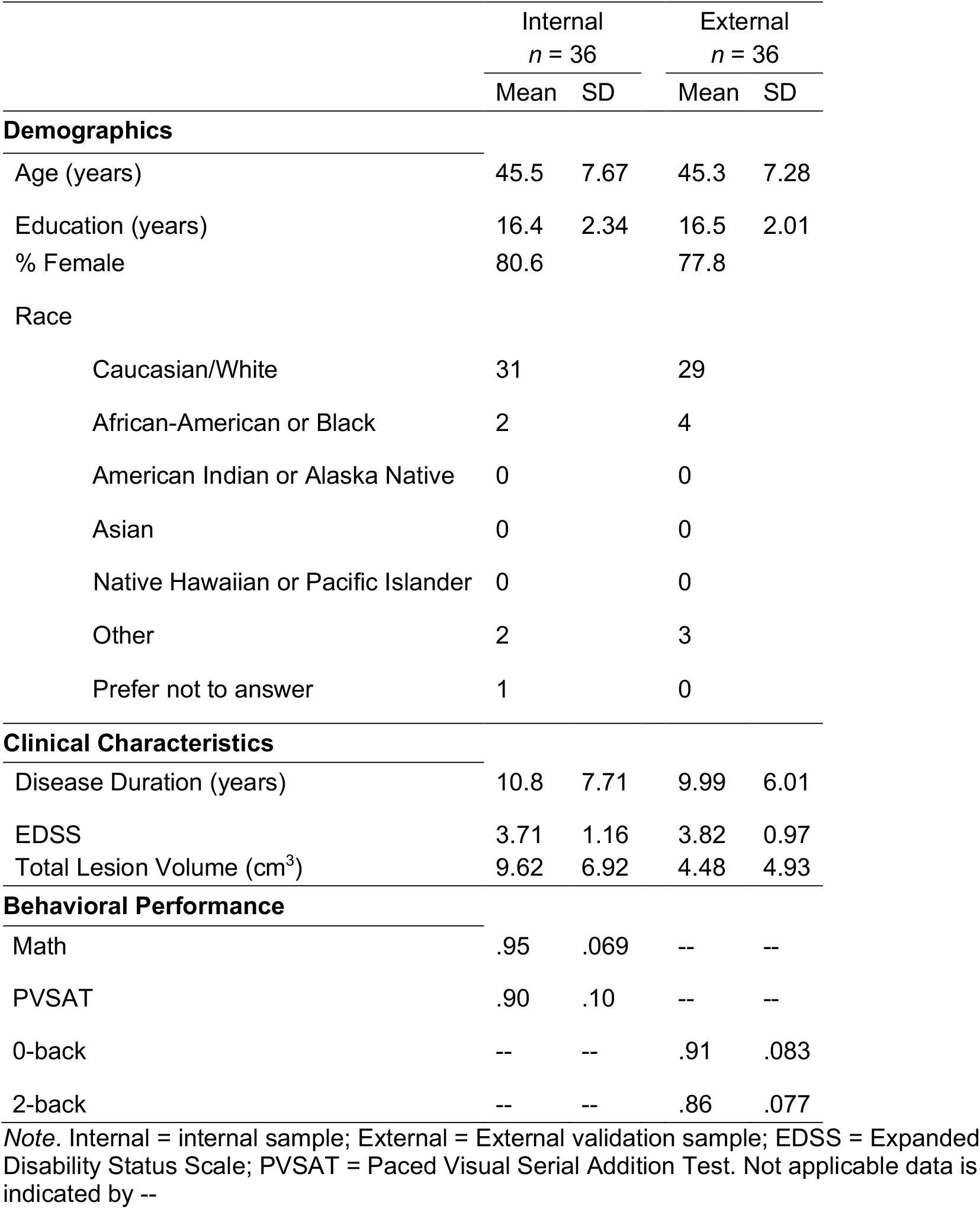
Demographics, clinical characteristics and behavioral performance of training and test samples

|  | Internal<br>*n* = 36 |  | External<br>*n* = 36 |  |
| --- | --- | --- | --- | --- |
|  | Mean | SD | Mean | SD |
| **Demographics** |  |  |  |  |
| Age (years) | 45.5 | 7.67 | 45.3 | 7.28 |
| Education (years) | 16.4 | 2.34 | 16.5 | 2.01 |
| % Female | 80.6 |  | 77.8 |  |
| Race |  |  |  |  |
| Caucasian/White | 31 |  | 29 |  |
| African-American or Black | 2 |  | 4 |  |
| American Indian or Alaska Native | 0 |  | 0 |  |
| Asian | 0 |  | 0 |  |
| Native Hawaiian or Pacific Islander | 0 |  | 0 |  |
| Other | 2 |  | 3 |  |
| Prefer not to answer | 1 |  | 0 |  |
| **Clinical Characteristics** |  |  |  |  |
| Disease Duration (years) | 10.8 | 7.71 | 9.99 | 6.01 |
| EDSS | 3.71 | 1.16 | 3.82 | 0.97 |
| Total Lesion Volume (cm<sup>3</sup>) | 9.62 | 6.92 | 4.48 | 4.93 |
| **Behavioral Performance** |  |  |  |  |
| Math | .95 | .069 | -- | -- |
| PVSAT | .90 | .10 | -- | -- |
| 0-back | -- | -- | .91 | .083 |
| 2-back | -- | -- | .86 | .077 |

*Note.* Internal = internal sample; External = External validation sample; EDSS = Expanded Disability Status Scale; PVSAT = Paced Visual Serial Addition Test. Not applicable data is indicated by --

### Network Construction

#### Internal Model Fitting

The results of network derivation using cross-validation revealed significant associations between observed performance and predictions based on the high (*r*_*s*_ = .47, permuted *p* = .011), low (*r*_*s*_ = .50, permuted *p* = .012), and full MS-wmCPM networks (*r*_*s*_ = .47, permuted *p* = .006; Table 2). Figure 1 displays the correlations between predicted and observed scores for each network and the corresponding permuted null distributions used for significance testing. Note, prediction accuracy indices of MSE and *R*^2^ are not reported for internal validation as these metrics overestimate the model fit when using cross-validation (Poldrack *et al*., 2019). Using a 9-fold cross validation with linear regression over 1000 iterations also resulted in a significant correlation between observed and predicted scores for the full network (median *r*_*s*_ = .47, median permuted *p* = .004). A single round of LOOCV with ridge regression resulted in significant prediction from the full model (*r*_*s*_ = .39, *p* = .020). The model with 100 permutations of 9-fold CV using ridge regression was not significant (median *r*_*s*_ = .31, median permuted *p* = .066), suggesting that the linear model with LOOCV was overfitting to the training sample. As motion was associated with PVSAT accuracy (*r*_*s*_ = -.42, *p* = .010), we include analyses controlling for the effects of motion on model performance in the Supplementary materials. All models, using linear regression and controlling for motion, were significant; models constructed using ridge regression were not significant.

**Table 2.** Model performance presented for each network

| Network Model | Sample | Network | $r_s$ | $p$ | MSE | prediction $R^2$ |
| --- | --- | --- | --- | --- | --- | --- |
| MS-wmCPM | Internal | High | .47 | .011<sup>a*</sup> |  |  |
|  |  | Low | .50 | .012<sup>a*</sup> |  |  |
|  |  | Full | .47 | .006<sup>1**</sup> |  |  |
| MS-wmCPM | External | High | .026 | .88 | .006 | -5.87% |
|  |  | Low | .084 | .63 | .006 | -3.78% |
|  |  | Full | -.047 | .78 | .006 | -5.13% |
| wmCPM | Internal | High | .20 | .24 | .010 | .63% |
|  |  | Low | -.33 | .046* | .009 | 14.4% |
|  |  | Full | .33 | .049* | .009 | 11.4% |
| wmCPM | External | High | .41 | .013* | .005 | 10.2% |
|  |  | Low | .51 | .002** | .005 | 15.5% |
|  |  | Full | .46 | .005** | .005 | 14.5% |

*Note.* The Spearman rank correlations between predicted and observed scores are presented here. The “Network Model” indicates which model was applied, to which “Sample” data, using which “Network”. WM = Working Memory; MS-wmCPM = working memory connectome-based predictive model derived in the internal sample of individuals with multiple sclerosis; wmCPM = working memory connectome-based predictive model derived in healthy young adults by Avery et al., (2019). <sup>a</sup> indicates permuted $p$ . \* = $p < .05$ , \*\* = $p < .01$ , \*\*\* = $p < .001$

**Figure 1.**
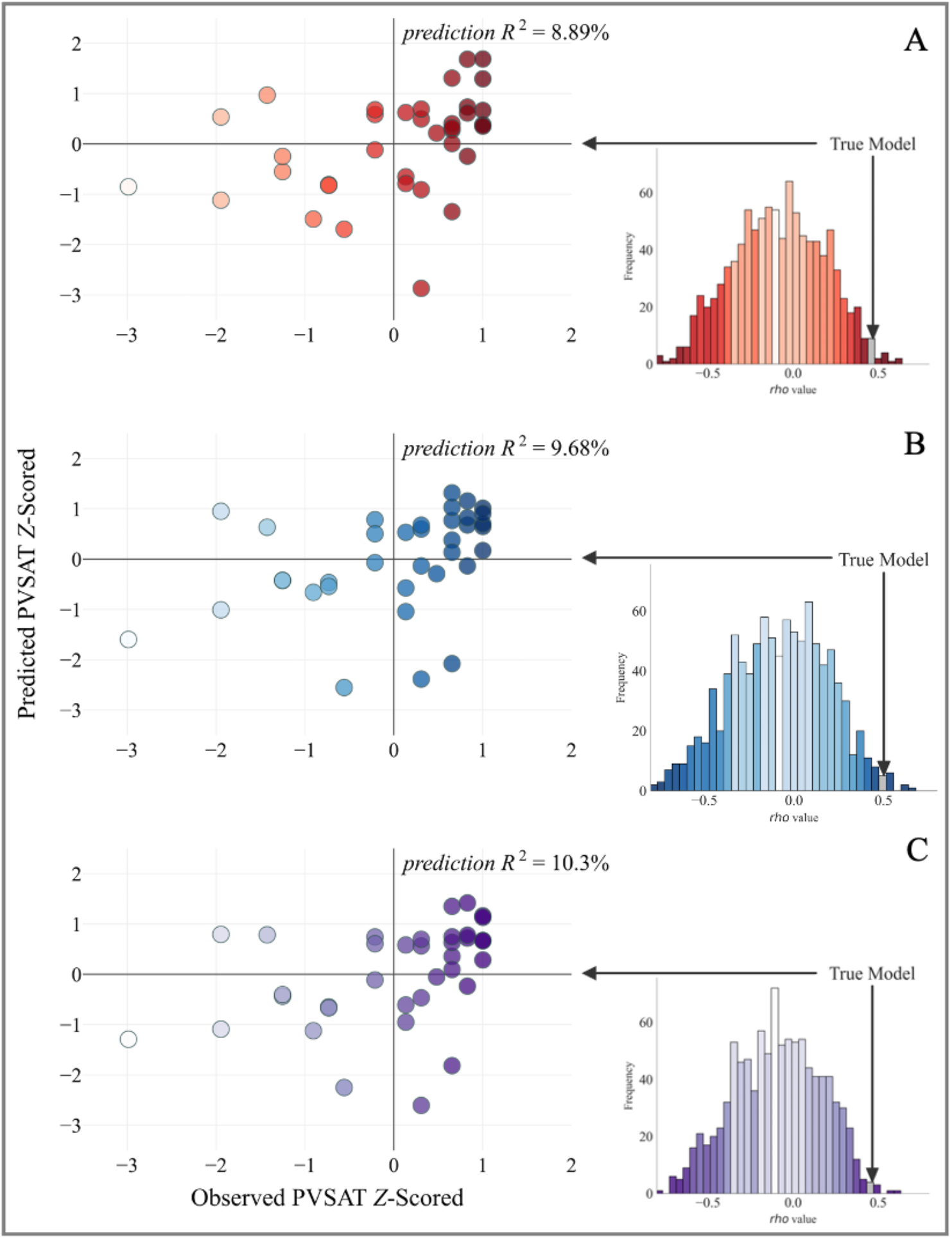
Internal model fit with permutation testing in the derivation sample. On the left are *z*-standardized PVSAT scores plotted observed against predicted from the A) high, B) low, and C) full MS-wmCPM networks. Annotations on each scatterplot represent *prediction R*^*2*^. Each scatterplot is paired with its respective null distribution of 1000 correlations between observed and predicted PVSAT scores from the CPM procedure with randomly-shuffled brain-behavior pairings. The gray bar depicts where the Spearman’s rank correlation between predicted and observed scores for the true model with actual brain-behavior pairings lies. Scatterplots and histograms were created using the python matplotlib package.

### External Validation

The final networks, containing edges that appeared in every round of cross validation, contained 189 edges in the high-WM network and 214 edges in the low-WM network (Figure 2). When testing the generalizability to the external validation sample, the MS-wmCPM did not successfully predict N-back performance in novel PwMS (Table 2). This model did not account for significant variance in observed N-back performance based on the high (*r*_*s*_ = .026; *p* = .88, MSE = .006, R^2^ = -5.87%), low (*r*_*s*_ = .084; *p* = .63, MSE = .006, R^2^ = -3.78%), or full MS-wmCPM network (*r*_*s*_ = -.047; *p* = .78, MSE = .006, R^2^ = -5.13%).

**Figure 2.**
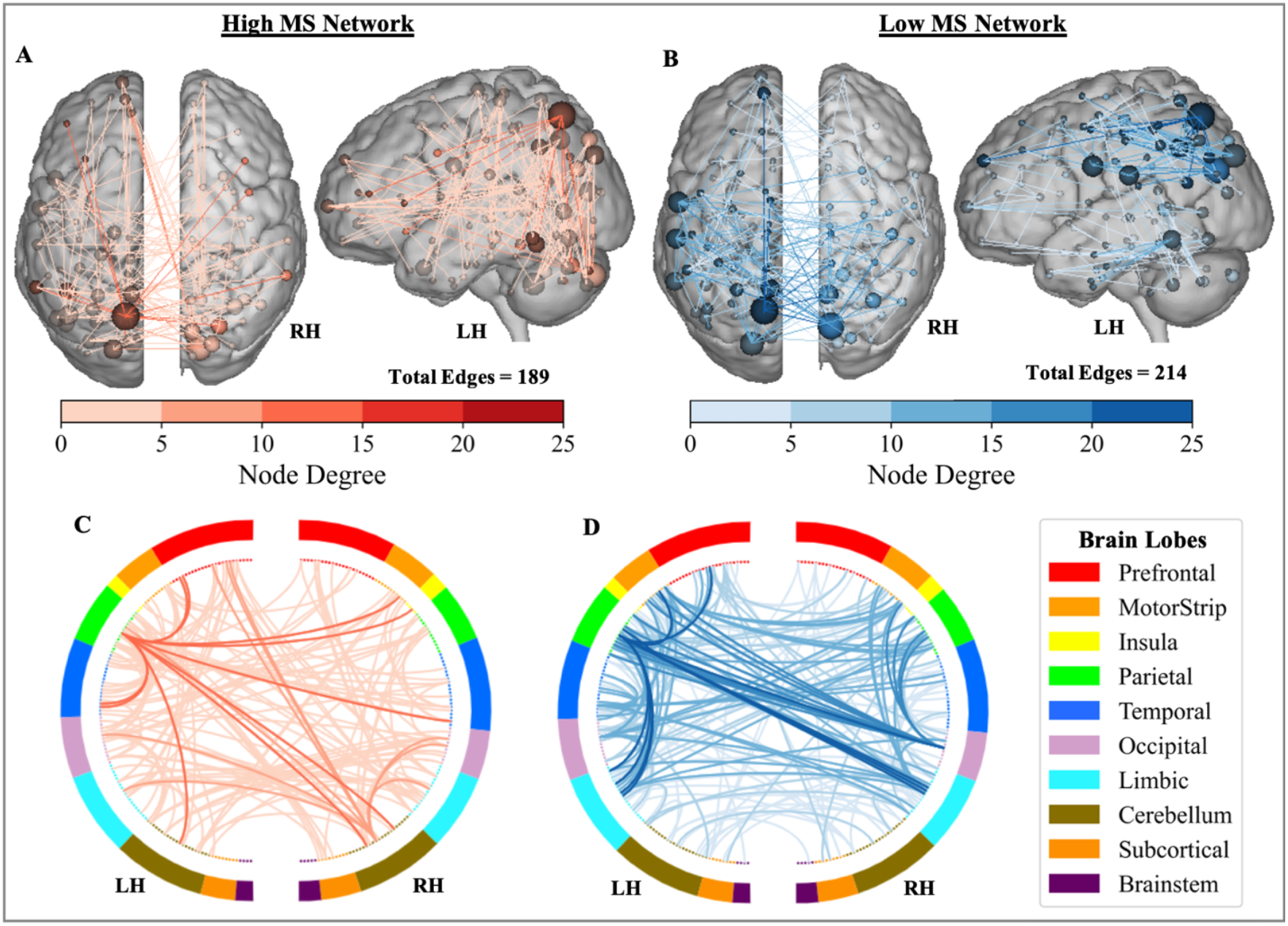
Distribution of the MS-wmCPM Networks. Edges of the high (red) and low (blue) working memory networks. Within the brains, nodes (spheres) represent regions and edges (lines) represent functional connections between nodes. Node size and edge color is proportionate to degree (i.e., number of edges). Ring plots depict the distribution of edges across macroscale regions of the brain. Darker lines in the ring plots indicate higher degree between brain lobes and match the color scale. Brain images and ring plots were created using the Yale BioImage Suite Connectivity Viewer Version 1.0.0 (https://bioimagesuiteweb.github.io/webapp/connviewer.html).

### Comparing Predictive Ability

The healthy wmCPM model (Avery *et al*., 2019) successfully predicted WM performance in both samples of PwMS. In the internal sample, models predicted significant variance from the low (*r*_*s*_ = -.33, *p* = .046, MSE = .009, R^2^ = 14.4%) and full networks (*r*_*s*_ = .33 *p* = .049, MSE = .009, R^2^ = 11.4%), but not the high network (*r*_*s*_ = .20, *p* = .24, MSE = .010, R^2^ = .63%). In the external sample, successful predictions of WM were found from all three wmCPM networks: high (*r*_*s*_ = .41, *p* = .013, MSE = .005, R^2^ = 10.2%), low (*r*_*s*_ = .51, *p* = .002, MSE = .005, R^2^ = 15.5%), and full (*r*_*s*_ = .46, *p* = .005, MSE = .005, R^2^ = 14.5%). Comparing model performance between the wmCPM and the MS-wmCPM when applied to the external validation sample, we found a statistically significant difference between the high (*z* = 1.97, *p* =.049), low (*z* = 2.55, *p* =.011), and full networks (*z* = 2.87, *p* =.004), such that the healthy wmCPM outperformed all three of the MS-wmCPM models. Model performance was not compared between the healthy and MS networks for the internal sample as in-sample model fit estimates likely inflate prediction accuracy.

### Functional Anatomy of WM Networks

Comparing the MS-wmCPM and wmCPM networks, the first striking difference is the smaller number of edges in the MS than healthy networks: 189 edges in high-MS vs 1674 edges in high-healthy, 214 edges in low-MS vs 1203 edges in low-healthy. It is plausible that successful external validation of only the wmCPM may in part be due to the larger raw number of edges comprising these networks. In addition, we found minimal overlap between the networks. Only 17 edges (8.99%) of the high MS-wmCPM were shared with the high wmCPM, and only 6 edges (2.80%) overlapped between the low MS-wmCPM and wmCPM networks.

We next examined the involvement of ten functionally defined canonical networks in each network, albeit interpreted with caution given the smaller size of the MS networks (Figure 3). We used a contribution index that accounts for network size and the total possible number of connections. For both high- and low-WM, MS networks showed greater involvement of visual nodes as compared to the healthy network. Examined separately, the high and low, and MS vs healthy networks illustrate observable differences involving the frontoparietal and default mode canonical networks (Figure 4).

**Figure 3.**
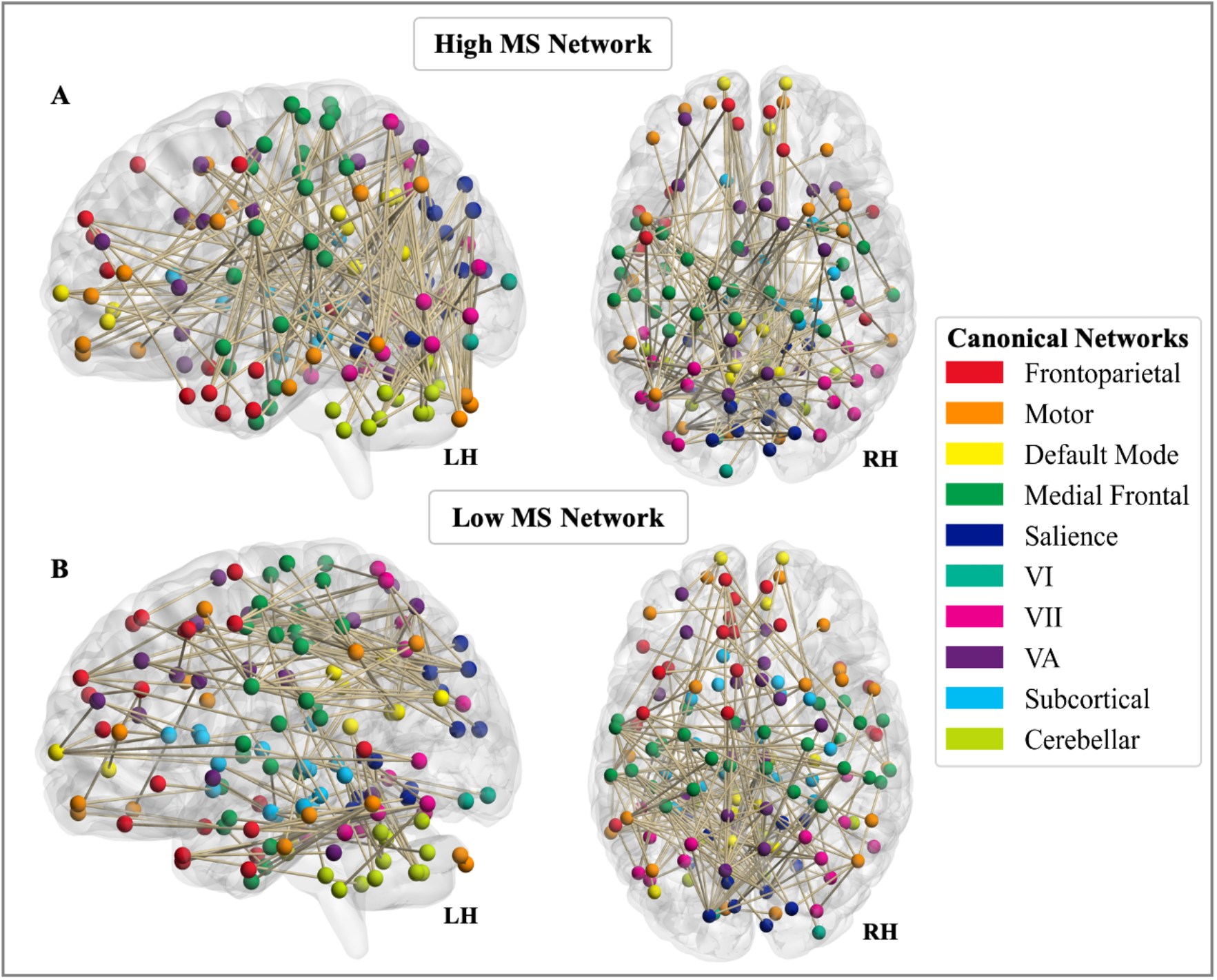
Involvement of canonical networks in MS-wmCPM. Functional anatomy of the A) high and B) low working memory networks derived in multiple sclerosis represented by involvement of ten canonical networks. Brain images were created using BrainNet Viewer version 1.7 (Xia *et al*., 2013).

**Figure 4.**
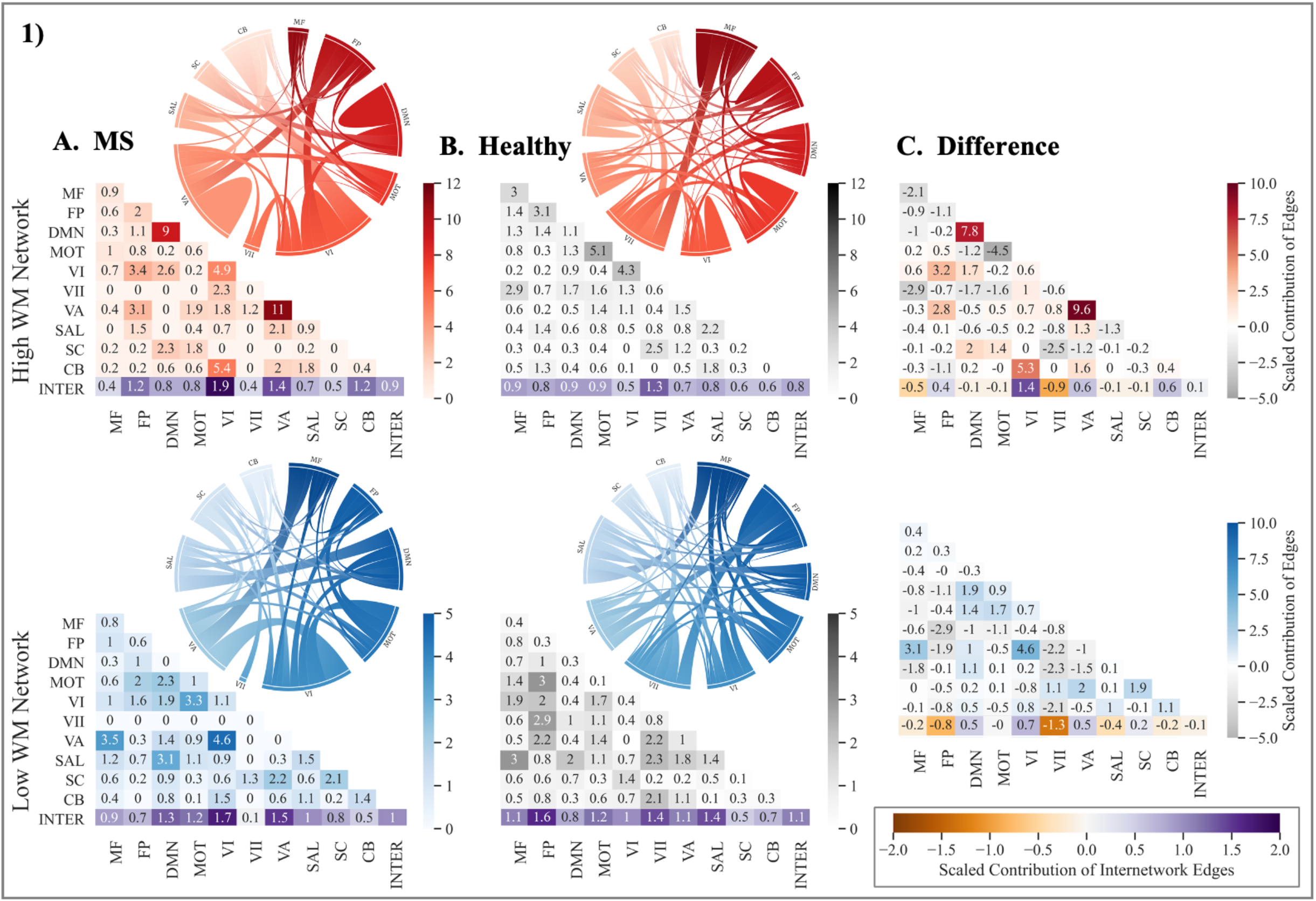
Contribution of canonical networks to working memory CPMs. The 1) high and 2) low working memory networks derived in A.) multiple sclerosis, B.) healthy adults, and 3) their difference (MS-Healthy). Degree of contribution is represented by color opacity; for the third column representing difference, positive (red or blue) values indicate greater contribution in multiple sclerosis networks and negative (gray) values indicate a greater contribution in healthy networks. Diagonals represent intranetwork contribution and the bottom row depicts the average internetwork contribution. The average of internetwork contribution is presented as the last value (at the intersection of Inter-Inter in the matrices). Value > 1 indicates disproportionately large contribution. Note: MS = multiple sclerosis; WM = working memory; MF = Medial Frontal; FP = Frontoparietal; DMN = Default Mode Network; MOT = Motor; VI = Visual I; VII = Visual II; VA = Visual Association; SAL = Salience, SC = Subcortical; CB = Cerebellar; INTER = Internetwork. Matrices were generated using the seaborn python package, and ring plots were constructed using Flourish (https://flourish.studio/).

In general, the expected pattern of intranetwork and internetwork involvement was evident in the healthy networks. Greater intranetwork contribution of the frontoparietal network was prominent in the high network (related to better performance), while greater internetwork connections between the default mode network and the rest of the brain was evident in the low network (related to poorer performance). In contrast, the MS networks showed a reverse pattern. Compared to the high healthy network, the high MS network involved greater inter-frontoparietal network edges, suggesting that to perform better on a demanding WM task, PwMS may rely on greater integration between frontoparietal nodes and the rest of the brain. Additionally, intranetwork connectivity within the default mode was also greater in the high MS relative to the high healthy network. In line with this, the low MS networks involved greater internetwork contribution of the default mode, suggesting that reduced segregation of the default mode network may be related to poorer WM performance. Taken together, these comparisons illustrate that for better WM performance, compared to healthy individuals, PwMS may rely on greater intra-default mode network connectivity and inter-frontoparietal network connectivity with diffuse functional networks.

## Discussion

This study aimed to apply an advanced predictive modeling approach to identify a functional network predictive of WM in PwMS. Using task-fMRI, we derived a predictive model of WM, and tested its generalizability to an independent sample of PwMS during completion of an alternative WM task. Although, the WM model generated using linear regression yielded significant predictions in the internal sample, it did not successfully generalize to a novel sample. Additionally, the WM model built using ridge regression was not significant, suggesting that the linear regression procedure may have been overfitting to the training sample. In contrast, WM networks derived in healthy young adults (Avery *et al*., 2019) successfully predicted WM performance in both samples of PwMS. These findings complement Avery and colleagues’ (2019) external validation of these healthy WM networks to clinical populations with varying degrees of cognitive impairment. The healthy and MS networks were found to have minimal overlap in edges and differences in involvement of canonical networks. These findings suggest that WM networks derived in healthy adults may demonstrate more robust, generalizable predictions of WM across heterogeneous clinical samples.

The challenge of identifying a reliable neuromarker of cognitive dysfunction in MS has persisted for decades against a backdrop of growing evidence that cognitive function in MS depends on a diffuse functional tapestry (Manca *et al*., 2018). The WM network derived in MS in this study, spanning all 10 canonical networks, suggests that WM is indeed an orchestration of widespread areas of the brain. The distribution of this network across disparate parts of the brain aligns with previous studies in healthy adults suggesting that WM calls upon distributed patterns of functional circuitry (Gazzaley *et al*., 2004; D’Esposito, 2007; Breukelaar *et al*., 2018; Eryilmaz *et al*., 2020). Further, this expands upon correlational research in MS demonstrating associations between WM performance and greater recruitment of broad regions during task-based MRI relative to healthy controls (Mainero *et al*., 2004; Wishart *et al*., 2004).

Contrary to our hypothesis, the model did not accurately predict WM function in a novel sample of PwMS. Connectome-based predictive modeling has been successful in predicting many different measures across clinical populations with heterogeneous presentations and symptoms. Specifically, CPM has been used to predict symptom improvement after pharmacotherapy in major depressive disorder (Ju *et al*., 2020), waist circumference and fasting insulin in overweight and obese individuals (Farruggia *et al*., 2020), global cognition in samples of individuals with mild cognitive impairment and Alzheimer’s disease (Lin *et al*., 2018), and self-reported executive dysfunction and memory in patients with breast cancer (Henneghan *et al*., 2020). Of note, although some of these studies included repeated, longitudinal assessment of networks (Henneghan *et al*., 2020; Ju *et al*., 2020), none of these studies in clinical populations included independent samples to test the generalizability of identified networks. Thus, successful prediction of WM in our derivation sample is consistent with prior clinical work but our results illuminate potential limitations to generalizability. Given the success of WM networks derived in young adults to generalize to clinical samples of older adults with varying levels of memory impairment (Avery *et al*., 2019), we also believe that failure of the WM model to generalize to an independent dataset of MS is not attributable to some central limitation of the method used. However, alternative methods to linear regression (i.e., ridge/lasso regression) may be better suited for small clinical samples as they account for multicollinearity in data with a large set of features (e.g., edges) and counter overfitting to the heterogeneity in the training data. Our findings support the use of independent samples and penalizing machine learning methods for stringent tests of generalizability, particularly when aiming to identify robust networks for use as targets in intervention.

Another explanation for the unsuccessful generalization may be related to the inherent structural damage observed in MS. There is evidence that as early as one year from a single neurological episode, white matter damage precedes functional restructuring at both global and modular levels (Koubiyr *et al*., 2019). Some modeling research posits that functional connectivity changes follow an inverted-U curve such that white matter damage initially leads to higher local functional connectivity, followed by a subsequent decrease, suggesting that the topology and timing of structural damage are important determinants of functional connectivity patterns in MS (Tewarie *et al*., 2018). Relatedly, as the derivation sample in the present study had high variability in disease duration (*M* = 10.8, *SD* = 7.71 years), it may have been comprised of individuals at variable disease stages, and thus with heterogeneous structural and consequent functional connectivity profiles. Given that the final network selects functional connections that were strongly related to behavior in every individual in the derivation sample, it is plausible that heterogeneous reorganization of functional connections at the individual level excluded edges that were relevant in only some individuals. As such, structural disconnection and consequent functional reorganization may have been too variable across individuals for identification of a reliable signature of WM.

Interestingly, WM networks derived in healthy young adults demonstrated robust generalization to both datasets of PwMS. In general, both healthy and MS-derived high-WM networks were characterized by greater contributions of intranetwork connectivity, suggesting that greater functional integrity of canonical networks is predictive of better WM performance. In contrast, the low-WM networks consisted of greater contributions of internetwork connections suggesting that decreased segregation between canonical communities predicts worse WM performance.

Comparing the relative involvement of canonical networks in the MS versus healthy networks, we found several interesting, yet complementary differences. The high MS network involved greater connections between the frontoparietal network and other networks, suggesting that functional integration may support better working memory performance. Our results also show that the high-WM network in MS involved greater connections within the default mode network while the low-WM network had a greater contribution of connections between the default mode network and other network nodes. This is consistent with evidence that even in early stages of disease, PwMS show increased functional connectivity in the default mode network, which has been posited as a compensatory mechanism to maintain cognitive efficiency in the presence of structural damage (Louapre *et al*., 2014). Interestingly, these patterns are the reverse of those observed in the healthy working memory networks where greater segregation of frontoparietal networks was related to better performance, whereas poorer segregation of the default mode network was linked to worse performance. These patterns illustrate that although the frontoparietal and default mode networks are of relevance, the segregation and integration of these networks may have differing contributions to performance in healthy individuals and PwMS.

Several important limitations require consideration. First, our sample sizes may have been underpowered to identify a robust neuromarker of WM. Recent studies suggest that samples on the order of ≥ 100 participants may be necessary for deriving reliable neural signatures of behavior (Poldrack *et al*., 2019). As the samples examined here were used as part of parent investigations of physical activity, our study was also restricted to individuals with relapsing remitting MS with mild disease severity and minimal cognitive dysfunction. There was also a significant difference in lesion volume between the two MS samples which may have contributed to differences in functional network topology and poor generalizability. This study may also have been limited by the use of linear regression for WM model derivation. This method was selected *a priori* to match the CPM procedure used in the healthy sample (Avery *et al*., 2019). In an exploratory analysis, we were unable to derive a significant model using ridge regression, suggesting that our MS-wmCPM may have been overfit to the internal sample and thus failed to generalize to the external sample of MS. Future clinical studies should leverage machine learning methods (i.e., ridge/lasso regression) to minimize the influence of training data and identify models with greater generalizability. To further enhance the predictive power and generalizability to independent samples of MS, researchers should seek to incorporate structural connectivity and/or lesion pathology data. Finally, a future direction may be to derive and compare models of WM derived in PwMS and healthy controls matched in age, gender, and education, to control for these factors and disentangle differences between healthy and clinical connectomes.

## Conclusion

This study demonstrates the utility of connectome-based predictive models in predicting WM function in PwMS. Models derived in healthy individuals outperformed models derived in MS, suggesting translational potential for use in clinical populations. Future well-powered investigations could build upon these findings by carefully sampling individuals with MS along important parameters including disease duration, disease subtypes, and levels of cognitive preservation/impairment. Moreover, this literature would benefit from continued use of predictive and whole-brain search light approaches to understand potential brain mechanisms of cognitive functions in MS. Joint structure-function connectomes from larger samples provide promising avenues to better understand the complex neural systems supporting cognition and identify reliable targets for intervention in MS.

## Acknowledgements

We are grateful for our study participants. We also thank Alisha Janssen, Ph.D., Brittney Schirda, Ph.D., Patrick Whitmoyer, Ph.D., Elizabeth Herring, Michael McKenna, Jacqueline Levine, Lauren Cea, Daniel Evans, Beth Patterson, Megan Fisher, Siobhan McDermott, Clara Huffman, and Alisha Bhagwat for instrumental support in recruitment and data collection. Finally, we express immense gratitude to Emily Avery and Monica D. Rosenberg, Ph.D., for providing access to the healthy working memory CPM networks used in the current study.

## Authors’ Contributions

H.R.M. contributed to conceptualization, data curation, formal analysis, investigation, methodology, software, writing – original draft, visualization, project administration.

S.FZ. contributed to validation, formal analysis, conceptualization, methodology, software, writing – review & editing, visualization.

A.S. contributed to data curation, software, writing – review & editing.

J.A.N. contributed to conceptualization, supervision, writing – review & editing.

R.S.P. contributed to conceptualization, supervision, funding acquisition, resources, writing – review & editing.

## Citation Diversity Statement

Papers by women and other minorities are under-cited relative to the overall number in the field (Maliniak *et al*., 2013; Mitchell *et al*., 2013; Caplar *et al*., 2017; Chakravartty *et al*., 2018; Dion *et al*., 2018, Dworkin *et al*., 2020*a, b*). This evidence of bias in citation practices prompted the current citation diversity analysis conducted on the main text and Supplementary Materials. After proactively choosing references that reflect diversity in thought, we used an open source automatic binary classification of gender and race based on the first names of the first and last authors (Ambekar *et al*., 2009; Sood and Laohaprapanon, 2018, Dworkin *et al*., 2020*b*; Zhou *et al*., 2020). Excluding self-citations to the first and last authors of the current study, our references contain 11.93% woman(first)/woman(last), 22.0% man/woman, 25.61% woman/man, and 40.47% man/man. Additionally, our references contain 13.16% author of color (first)/author of color(last), 13.82% white author/author of color, 20.31% author of color/white author, and 52.7% white author/white author. These methods are limited in that: (i) names, pronouns, social media and Wikipedia profiles, and Census entries used to construct the databases may not be indicative of gender or racial/ethnic identity, (ii) they do not account for intersex, non-binary, transgender, Indigenous and mixed-race authors, or those who may face differential biases due to the ambiguous racialization or ethnicization of their names. This is a start, and we encourage future work to support equitable practices in science.

## References

Ambekar A, Ward C, Mohammed J, Male S, Skiena S. Name-ethnicity classification from open sources. In: In Proceedings of the 15th ACM SIGKDD international conference on Knowledge Discovery and Data Mining. 2009. p. 49–58

Avery EW, Yoo K, Rosenberg MD, Greene AS, Gao S, Na DL, et al. Distributed Patterns of Functional Connectivity Predict Working Memory Performance in Novel Healthy and Memory-impaired Individuals. J Cogn Neurosci 2019: 1–15.

Barch DM, Burgess GC, Harms MP, Petersen SE, Schlaggar BL, Corbetta M, et al. Function in the Human Connectome: Task-fMRI and Individual Differences in Behavior. NeuroImage 2013; 80: 169–89.

Barron DS, Gao S, Dadashkarimi J, Greene AS, Spann MN, Noble S, et al. Task-Based Functional Connectomes Predict Cognitive Phenotypes Across Psychiatric Disease [Internet]. bioRxiv 2019[cited 2019 Jul 3] Available from: http://biorxiv.org/lookup/doi/10.1101/638825

Beck AT, Steer RA, Brown GK. Manual for the Beck Depression Inventory-II. San Antonio, TX: Psychological Corporation; 1996

Breukelaar IA, Williams LM, Antees C, Grieve SM, Foster SL, Gomes L, et al. Cognitive ability is associated with changes in the functional organization of the cognitive control brain network. Hum Brain Mapp 2018; 39: 5028–38.

Cabeza R, Nyberg L, Park D C. Cognitive Neuroscience of Aging: Linking Cognitive And Cerebral Aging. Oxford University Press; 2016

Caplar N, Tacchella S, Birrer S. Quantitative Evaluation of Gender Bias in Astronomical Publications from Citation Counts. Nat Astron 2017; 1: 0141.

Chakravartty P, Kuo R, Grubbs V, McIlwain C. #CommunicationSoWhite. J Commun 2018; 68: 254–66.

Chiaravalloti N, Hillary FG, Ricker JH, Christodoulou C, Kalnin AJ, Liu WC, et al. Cerebral activation patterns during working memory performance in multiple sclerosis using fMRI. J Clin Exp Neuropsychol 2005; 27: 33–54.

Chiaravalloti ND, DeLuca J. Cognitive impairment in multiple sclerosis. Lancet Neurol 2008; 7: 1139–51.

Colorado RA, Shukla K, Zhou Y, Wolinsky JS, Narayana PA. Multi-task functional MRI in multiple sclerosis patients without clinical disability. NeuroImage 2012; 59: 573–81.

Cui Z, Gong G. The effect of machine learning regression algorithms and sample size on individualized behavioral prediction with functional connectivity features. NeuroImage 2018; 178: 622–37.

D’Esposito M. From cognitive to neural models of working memory. Philos Trans R Soc B Biol Sci 2007; 362: 761–72.

Diedenhofen B, Musch J. cocor: A Comprehensive Solution for the Statistical Comparison of Correlations. PLOS ONE 2015; 10: e0121945.

Dion ML, Sumner JL, Mitchell SM. Gendered Citation Patterns across Political Science and Social Science Methodology Fields. Polit Anal 2018; 26: 312–27.

Dunn OJ, Clark V. Comparison of Tests of the Equality of Dependent Correlation Coefficients. J Am Stat Assoc 1971; 66: 904–8.

Dworkin J, Zurn P, Bassett DS. (In)citing Action to Realize an Equitable Future. Neuron 2020; 106: 890–4.

Dworkin JD, Linn KA, Teich EG, Zurn P, Shinohara RT, Bassett DS. The extent and drivers of gender imbalance in neuroscience reference lists [Internet]. Nat Neurosci 2020[cited 2020 Jun 28] Available from: http://www.nature.com/articles/s41593-020-0658-y

Eryilmaz H, Dowling KF, Hughes DE, Rodriguez-Thompson A, Tanner A, Huntington C, et al. Working memory load-dependent changes in cortical network connectivity estimated by machine learning. NeuroImage 2020; 217: 116895.

Farruggia MC, van Kooten MJ, Perszyk EE, Burke MV, Scheinost D, Constable RT, et al. Identification of a brain fingerprint for overweight and obesity. Physiol Behav 2020; 222: 112940.

Finn ES, Shen X, Scheinost D, Rosenberg MD, Huang J, Chun MM, et al. Functional connectome fingerprinting: identifying individuals using patterns of brain connectivity. Nat Neurosci 2015; 18: 1664–71.

Folstein MF, Folstein SE, McHugh PR. ‘Mini-mental state’. A practical method for grading the cognitive state of patients for the clinician. J Psychiatr Res 1975; 12: 189–98.

Forn C, Barros-Loscertales A, Escudero J, Benlloch V, Campos S, Parcet MA, et al. Compensatory activations in patients with multiple sclerosis during preserved performance on the auditory N-back task. Hum Brain Mapp 2007; 28: 424–30.

Fos LA, Greve KW, South MB, Mathias C, Benefield H. Paced Visual Serial Addition Test: An Alternative Measure of Information Processing Speed. Appl Neuropsychol 2000; 7: 140–6.

Fountain-Zaragoza S, Samimy S, Rosenberg MD, Prakash RS. Connectome-based models predict attentional control in aging adults. NeuroImage 2019; 186: 1–13.

Gazzaley A, Rissman J, D’Esposito M. Functional connectivity during working memory maintenance. Cogn Affect Behav Neurosci 2004; 4: 580–99.

Giorgio A, Stromillo ML, De Leucio A, Rossi F, Brandes I, Hakiki B, et al. Appraisal of Brain Connectivity in Radiologically Isolated Syndrome by Modeling Imaging Measures. J Neurosci 2015; 35: 550–8.

Greene AS, Gao S, Scheinost D, Constable RT. Task-induced brain state manipulation improves prediction of individual traits [Internet]. Nat Commun 2018; 9[cited 2018 Dec 7] Available from: http://www.nature.com/articles/s41467-018-04920-3

Henneghan AM, Gibbons C, Harrison RA, Edwards ML, Rao V, Blayney DW, et al. Predicting Patient Reported Outcomes of Cognitive Function Using Connectome-Based Predictive Modeling in Breast Cancer. Brain Topogr 2020; 33: 135–42.

Hillary FG, Chiaravalloti ND, Ricker JH, Steffener J, Bly BM, Lange G, et al. An Investigation of Working Memory Rehearsal in Multiple Sclerosis Using fMRI. J Clin Exp Neuropsychol 2003; 25: 965–78.

Hoerl AE, Kennard RW. Ridge Regression: Biased Estimation for Nonorthogonal Problems. Technometrics 2000; 42: 80–6.

Ju Y, Horien C, Chen W, Guo W, Lu X, Sun J, et al. Connectome-based models can predict early symptom improvement in major depressive disorder. J Affect Disord 2020; 273: 442–52.

Koubiyr I, Besson P, Deloire M, Charre-Morin J, Saubusse A, Tourdias T, et al. Dynamic modular-level alterations of structural-functional coupling in clinically isolated syndrome. Brain 2019; 142: 3428–39.

Kurtzke JF. Rating neurologic impairment in multiple sclerosis An expanded disability status scale (EDSS). Neurology 1983; 33: 1444–52.

Lin Q, Rosenberg MD, Yoo K, Hsu TW, O’Connell TP, Chun MM. Resting-State Functional Connectivity Predicts Cognitive Impairment Related to Alzheimer’s Disease [Internet]. Front Aging Neurosci 2018; 10[cited 2018 Dec 18] Available from: https://www.frontiersin.org/articles/10.3389/fnagi.2018.00094/full

Louapre C, Perlbarg V, García-Lorenzo D, Urbanski M, Benali H, Assouad R, et al. Brain networks disconnection in early multiple sclerosis cognitive deficits: An anatomofunctional study. Hum Brain Mapp 2014; 35: 4706–17.

Macías Islas M, Ciampi E. Assessment and Impact of Cognitive Impairment in Multiple Sclerosis: An Overview. Biomedicines 2019; 7: 22.

Mainero C, Caramia F, Pozzilli C, Pisani A, Pestalozza I, Borriello G, et al. fMRI evidence of brain reorganization during attention and memory tasks in multiple sclerosis. NeuroImage 2004; 21: 858–67.

Maliniak D, Powers R, Walter BF. The Gender Citation Gap in International Relations. Int Organ 2013; 67: 889–922.

Manca R, Sharrack B, Paling D, Wilkinson ID, Venneri A. Brain connectivity and cognitive processing speed in multiple sclerosis: A systematic review. J Neurol Sci 2018; 388: 115–27.

Meijer KA, Eijlers AJ, Douw L, Uitdehaag BM, Barkhof F, Geurts JJ, et al. Increased connectivity of hub networks and cognitive impairment in multiple sclerosis. Neurology 2017; 88: 2107–14.

Mitchell SM, Lange S, Brus H. Gendered Citation Patterns in International Relations Journals. Int Stud Perspect 2013; 14: 485–92.

Nicholas JA, Electricwala B, Lee LK, Johnson KM. Burden of relapsing-remitting multiple sclerosis on workers in the US: a cross-sectional analysis of survey data [Internet]. BMC Neurol 2019; 19[cited 2020 May 9] Available from: https://bmcneurol.biomedcentral.com/articles/10.1186/s12883-019-1495-z

Poldrack RA, Huckins G, Varoquaux G. Establishment of Best Practices for Evidence for Prediction: A Review [Internet]. JAMA Psychiatry 2019[cited 2020 Jan 6] Available from: https://jamanetwork.com/journals/jamapsychiatry/fullarticle/2756204

Rosenberg MD, Finn ES, Scheinost D, Papademetris X, Shen X, Constable RT, et al. A neuromarker of sustained attention from whole-brain functional connectivity. Nat Neurosci 2016; 19: 165–71.

Rosenberg MD, Hsu W-T, Scheinost D, Todd Constable R, Chun MM. Connectome-based Models Predict Separable Components of Attention in Novel Individuals. J Cogn Neurosci 2017; 30: 160–73.

Rovaris M, Filippi M, Minicucci L, Iannucci G, Santuccio G, Possa F, et al. Cortical/Subcortical Disease Burden and Cognitive Impairment in Patients with Multiple Sclerosis. 2000: 7.

Scheinost D, Noble S, Horien C, Greene AS, Lake EMR, Salehi M, et al. Ten simple rules for predictive modeling of individual differences in neuroimaging. NeuroImage 2019; 193: 35–45.

Shen X, Finn ES, Scheinost D, Rosenberg MD, Chun MM, Papademetris X, et al. Using connectome-based predictive modeling to predict individual behavior from brain connectivity. Nat Protoc 2017; 12: 506–18.

Shen X, Tokoglu F, Papademetris X, Constable T R. Groupwise whole-brain parcellation from resting-state fMRI data for network node identification. NeuroImage 2013; 82: 403–15.

Sood G, Laohaprapanon S. Predicting Race and Ethnicity From the Sequence of Characters in a Name [Internet]. ArXiv180502109 Stat 2018[cited 2020 Nov 5] Available from: http://arxiv.org/abs/1805.02109

Spielberg JM, McGlinchey RE, Milberg WP, Salat DH. Brain Network Disturbance Related to Posttraumatic Stress and Traumatic Brain Injury in Veterans. Biol Psychiatry 2015; 78: 210–6.

Sweet LH, Rao SM, Primeau M, Durgerian S, Cohen RA. Functional magnetic resonance imaging response to increased verbal working memory demands among patients with multiple sclerosis. Hum Brain Mapp 2006; 27: 28–36.

Tewarie P, Steenwijk MD, Brookes MJ, Uitdehaag BMJ, Geurts JJG, Stam CJ, et al. Explaining the heterogeneity of functional connectivity findings in multiple sclerosis: An empirically informed modeling study. Hum Brain Mapp 2018; 39: 2541–8.

Vacchi L, Rocca MA, Meani A, Rodegher M, Martinelli V, Comi G, et al. Working memory network dysfunction in relapse-onset multiple sclerosis phenotypes: A clinical-imaging evaluation. Mult Scler J 2017; 23: 577–87.

Van Essen D C, Ugurbil K, Auerbach E, Barch D, Behrens TEJ, Bucholz R, et al. The Human Connectome Project: A data acquisition perspective. NeuroImage 2012; 62: 2222–31.

Wallin MT, Culpepper WJ, Campbell JD, Nelson LM, Langer-Gould A, Marrie RA, et al. The prevalence of MS in the United States: A population-based estimate using health claims data. Neurology 2019; 92: e1029–40.

Wishart HA, Saykin AJ, McDonald BC, Mamourian AC, Flashman LA, Schuschu KR, et al. Brain activation patterns associated with working memory in relapsing-remitting MS. Neurology 2004; 62: 234–8.

Zhou D, Cornblath E J, Stiso, J, Teich EG, Dworkin JD, Blevins A S, et al. Gender Diversity Statement and Code Notebook v1.0 (Version v1.0) [Internet]. Zenodo 2020Available from: http://doi.org/10.5281/zenodo.3672110

