## Supplementary Materials for "Employing connectome-based models to predict working memory in multiple sclerosis"

**Imaging parameters**

Table 1

*Image Acquisition Parameters*

| | **Task** | | **Structural** | | **Lesion** | |
|---|---|---|---|---|---|---|
| Measure | PVSAT[a] | N-back[b] | MPRAGE[a] | MPRAGE[b] | Clear[a] | FLAIR[b] |
| Siemens 3T model | Trio | Prisma | Trio | Prisma | Trio | Prisma |
| Pulse sequence type | EPI | EPI | Gradient echo | Gradient echo | Fast spin-echo | Fast spin-echo |
| Parallel imaging parameters (band, type, acceleration factor) | Single, GRAPPA, 2 | Multiband, 3 | Single GRAPPA, 2 | Single | Single, GRAPPA 2 | Single, GRAPPA, 2 |
| Number of volumes | 408 | 910 (455 x 2 runs) | 1 | 1 | 1 | 1 |
| TR | 2000 ms | 1000 ms | 1950 ms | 1900 ms | 9000 ms | 9000 ms |
| TE | 30 ms | 28 ms | 4.44 ms | 4.44 ms | 67 ms | 95 ms |
| Flip angle | 73° | 50° | 12° | 12° | 120° | 135° |
| FOV | 220 mm | 240 mm | 256 mm | 256 mm | 256 mm | 256 mm |
| Acquisition matrix | 64 x 58 x 34 | 80 x 72 x 45 | 256 x 232 x 176 | 256 x 256 x 176 | 320 x 290 x 60 | 256 x 256 x 70 |
| Number of slices | 34 | 45 | 176 | 176 | 60 | 70s |
| Slice thickness | 2.5 mm | 3.0 mm | 1.0 mm | 1.0 mm | 2.0 mm | 2.0 mm |
| Voxel size | 3.4 $mm^3$ | 3.0 $mm^3$ | 1.0 $mm^3$ | 1.0 $mm^3$ | 0.8 mm x 0.8 mm x 2.0 mm | 1.0 mm x 1.0 mm x 2.0 mm |
| Bandwidth (Hz/Px) | 1594 | 2500 | 140 | 140 | 220 | 222 |
| Echo spacing | 0.71 ms | 0.5 ms | 10.1 ms | 10.1 ms | 9.52 ms | 8.65 ms |

| | | | | | | |
|---|---|---|---|---|---|---|
| Acquisition orientation | Axial-oblique | Axial-oblique | Sagittal | Sagittal | Transversal | Transversal |
| Acquisition order | Interleaved | Interleaved | Interleaved | Interleaved | Interleaved | Interleaved |

*Note.* [a] = internal validation sample; [b] = external validation sample; PVSAT = Paced Visual Serial Addition Test; MPRAGE = Magnetization Prepared Rapid Gradient Echo; FLAIR = Fluid Attenuated Inversion Recovery; EPI = Echo Planar Imaging; ms = millisecond; TR = repetition time; TE = echo time; FOV = field of view; Hx/Px = Hertz/Pixel.

**Data analysis**

Unless indicated otherwise, all processing steps were performed comparably for both samples using fMRIB Software Library version 5.0.9 (FSL; Smith *et al.*, 2004; Jenkinson *et al.*, 2012).

**Motion controls.** Participants with excessive head motion identified by both a mean framewise displacement > 0.2 mm (Power *et al.*, 2014) and > 10% of volumes with high motion (> 0.5 mm from previous volume) were excluded from analysis. Further control of motion artifacts was achieved by inclusion of 24 motion parameters (six translation and rotation realignment parameters, six temporal derivatives, and their squares), as well as volumes with > 0.5 mm framewise displacement from the previous (Power *et al.*, 2012) in nuisance regression.

**Lesion controls.** Identification and filling of lesions in the MPRAGE image was completed to improve registration and automated segmentation of nuisance regressors. For the internal validation sample, lesions were segmented from CLEAR images using both automated and manual procedures via JIM software (Xinapse Systems, www.xinapse.com). For the external validation sample, acquisition of high-quality FLAIR images permitted fully automated segmentation using the Lesion Segmentation Toolbox (LST; Schmidt *et al.*, 2012) (www.statistical-modelling.de/lst.html) version 2.0.15 for SPM. Thresholds for segmentation were determined separately in each sample using visual inspection. Using fMRIB's lesion filling tool (Battaglini *et al.*, 2012), the resulting lesion masks were applied to the MPRAGE image to fill lesion voxels with the intensity of nearby non-lesion voxels. The filled MPRAGE images were used as input for fMRIB's Automated Segmentation Tool (FAST; Zhang *et al.*, 2001) wherein white matter (WM) and cerebrospinal fluid (CSF) masks were created. The mean EPI signals within WM, CSF, and lesion masks attained after standard preprocessing were later entered as nuisance regressors.

**Image preprocessing.** The order of preprocessing steps was: motion correction using fMRIB's Linear Image Registration Tool (MCFLIRT; Jenkinson *et al.*, 2002), distortion correction using fMRIB's Utility for Geometrically Unwarping EPIs (Jenkinson *et al.*, 2012), interleaved slice timing correction for non-multiband data (the internal validation sample only), brain extraction of 4D data (BET; Smith, 2002), spatial smoothing using a 6mm full width at half maximum kernel, linear registration of 4D data to the participant's respective lesion-filled MPRAGE image, and nonlinear registration of 4D data to the standard MNI brain.

**Nuisance regression.** All processing steps requiring regression (i.e., temporal filtering, motion correction, nuisance regression) were subsequently completed in a single step, as supported by recent evidence from Lindquist and colleagues demonstrating that modular preprocessing (involving iterative regressions) reintroduces noise in studies of functional connectivity (Lindquist *et al.*, 2019). Thus, we conducted a single regression step that included high-pass temporal filtering (0.01 Hz) and regression of 24 motion parameters, high-motion confound volumes, and mean signal in WM, CSF, global signal and lesions. The 4D residuals from this regression were used in subsequent analyses.

**Effects of Motion**

Motion was significantly associated with PVSAT accuracy in the internal validation sample (mean frame-to-frame displacement = .12 mm; $\rho$ = -.42, $p$ = .010). Prior evidence indicates that motion during a WM task in people with multiple sclerosis is related to task difficulty and cognitive function, such that as task difficulty increases, those with lower cognitive functioning demonstrate greater motion (Wylie *et al.*, 2012). Given the overlapping variance between motion and cognitive ability, we did not control for motion at the edge selection step. In the internal validation sample, a partial Spearman correlation between predicted and observed scores controlling for motion resulted in non-significant correlations for all three networks: high ($r$ = .21, $p$ = .24), low ($r$ = .22; $p$ = .21), and full ($r$ = .22, $p$ = .20), suggesting overlapping variance between motion and WM performance. The correlation between 2-back accuracy and motion was not significant in the external validation sample ($r$ = .12, $p$ = .30).
